## Supplementary Figure for "Systematically developing a registry of splice-site creating variants utilizing massive publicly available transcriptome sequence data"

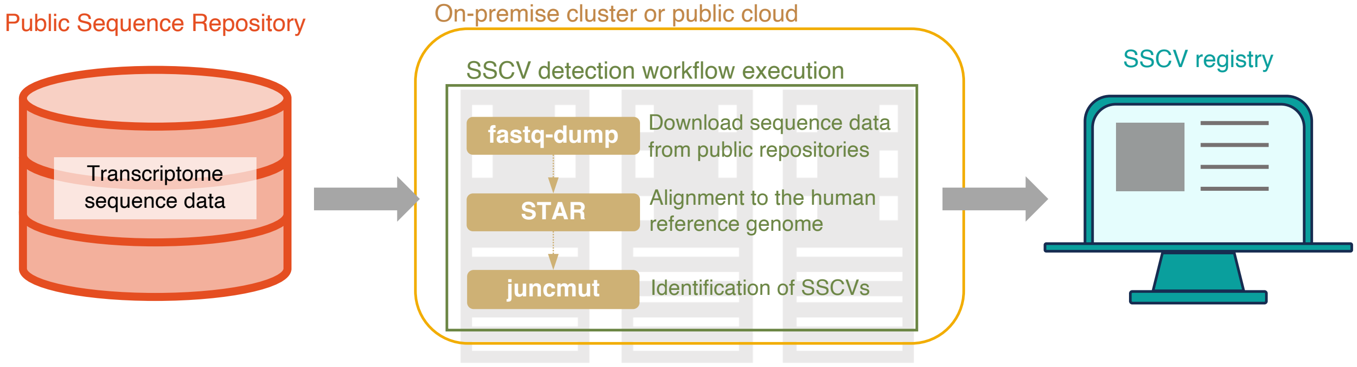


**Supplementary Figure 1: Overview of the proposed framework for detecting splice-site creating variants from raw sequencing data registered in Sequence Read Archive.** The downloaded sequence data is processed in an on-premises or public cloud computing environment and identified SSCVs are transferred to the SSCV registry and provided to the community.

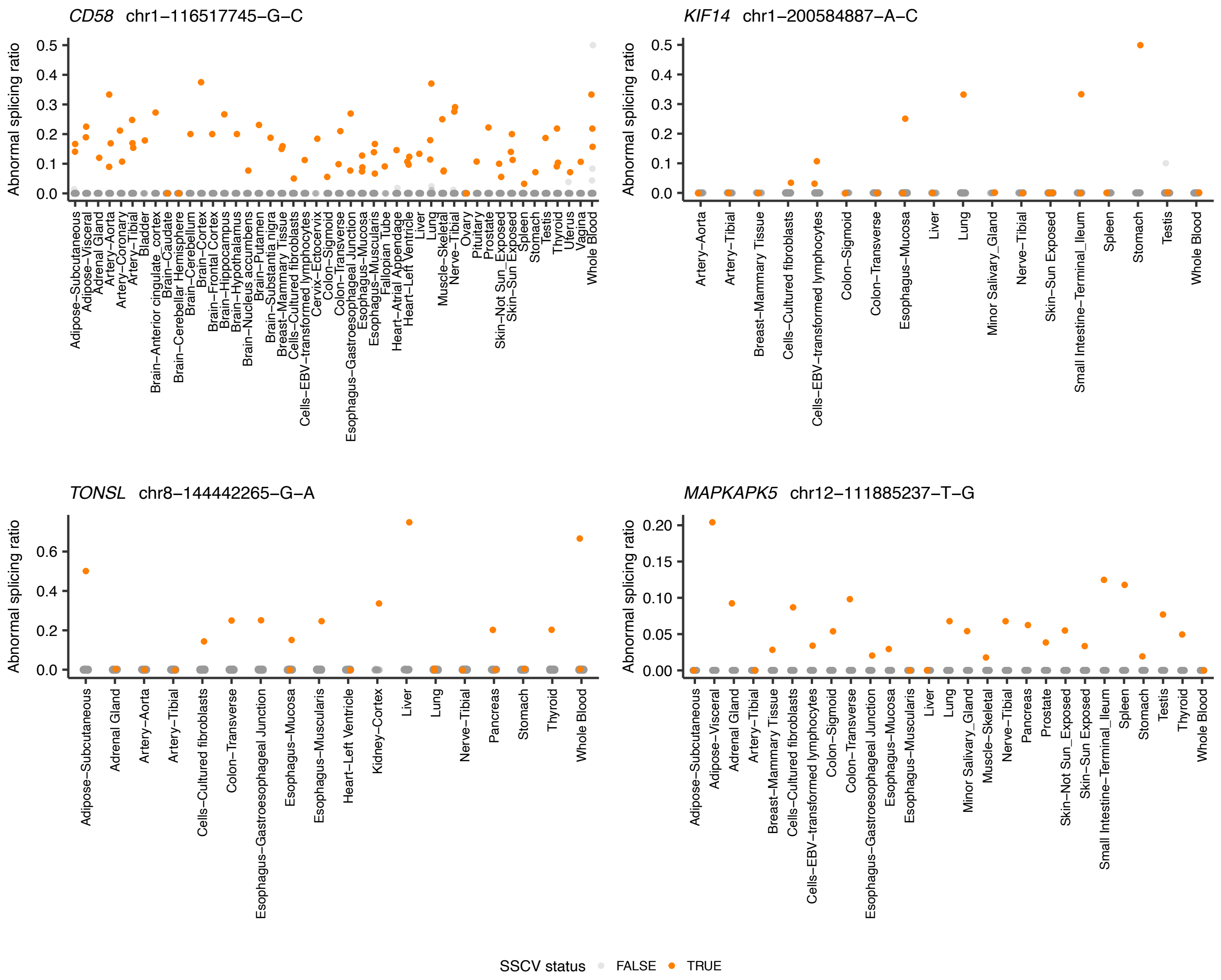


**Supplementary Figure 2: Relative ratios of corresponding abnormal splicing ratio for samples with and without the SSCVs across tissues measured using GTEx transcriptome data.** Each point shows an individual colored by SSCV mutation status. P-values, calculated using the one-sided Wilcoxon rank sum test to measure differences in abnormal splicing ratios between samples with and without SSCVs for each tissue, were integrated by Fisher's method. These values were $5.21\times10^{-120}$ in the top right and below $1.00\times10^{-320}$ in the other tissues. See also Figure 1f.

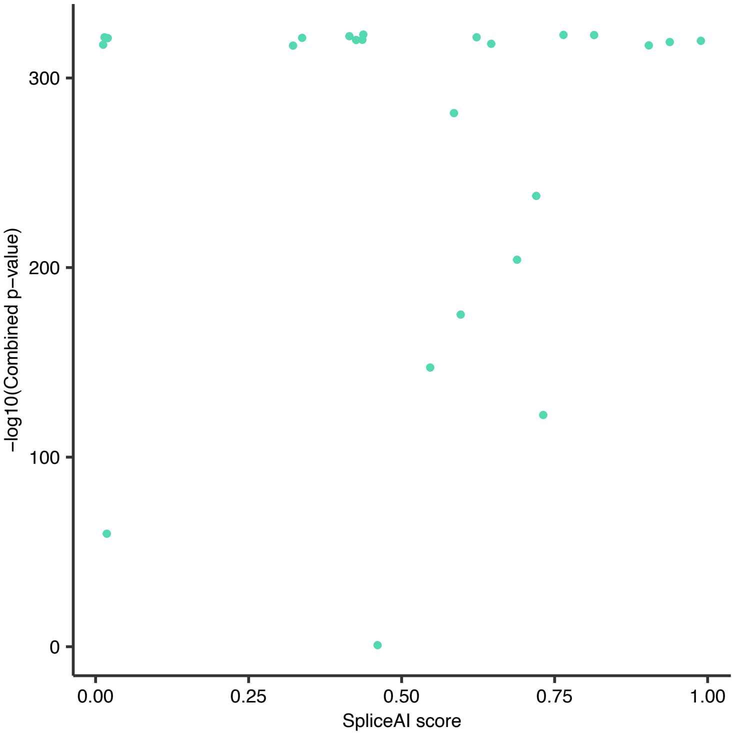


**Supplementary Figure 3: Scatter plot illustrating the relationship between the tissue-wide combined p-values for abnormal splicing ratio and the SpliceAI scores.** Note that the points are slightly jittered both horizontally and vertically to enhance visibility. Most detected SSCVs tend to have higher SpliceAI scores. However, some variants with a SpliceAI score of zero (specifically below 0.1, which is not displayed in the annotation program) still exhibit a high combined p-value.

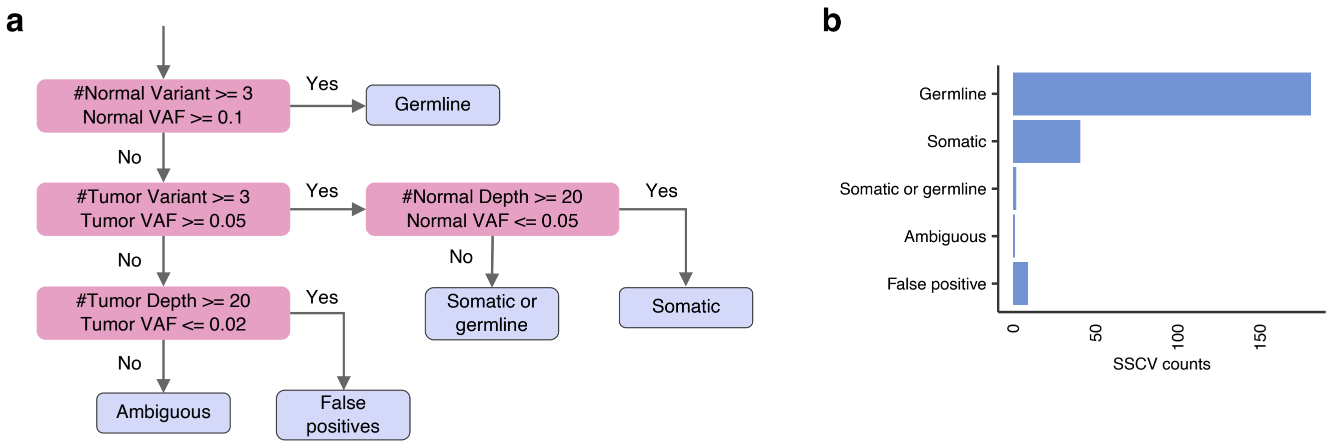


**Supplementary Figure 4: Assessment of juncmut approach using TCGA transcriptome and exome sequencing data.** (a) A flowchart illustrating the classification of genome-level mutation statuses for SSCVs. Classification is based on sequencing depth, variant count (number of SSCV containing reads), and variant allele frequency (VAF, ratio of SSCV containing reads to the total reads covering the position of SSCVs). Statuses are categorized as “germline,” “somatic,” “somatic or germline,” “ambiguous”, and “false positive.” (b) The number of SSCVs categorized by their inferred mutation status.

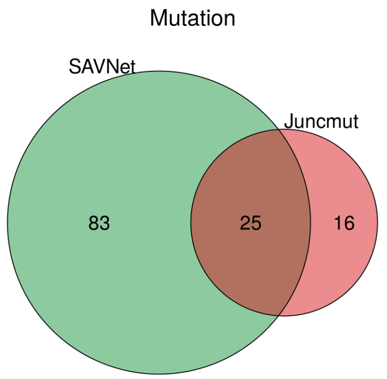


**Supplementary Figure 5: Venn diagrams showing the overlap of SSCVs identified by juncmut and the previous approach (SAVNet).** Refer to the Methods section for details regarding the comparison method.

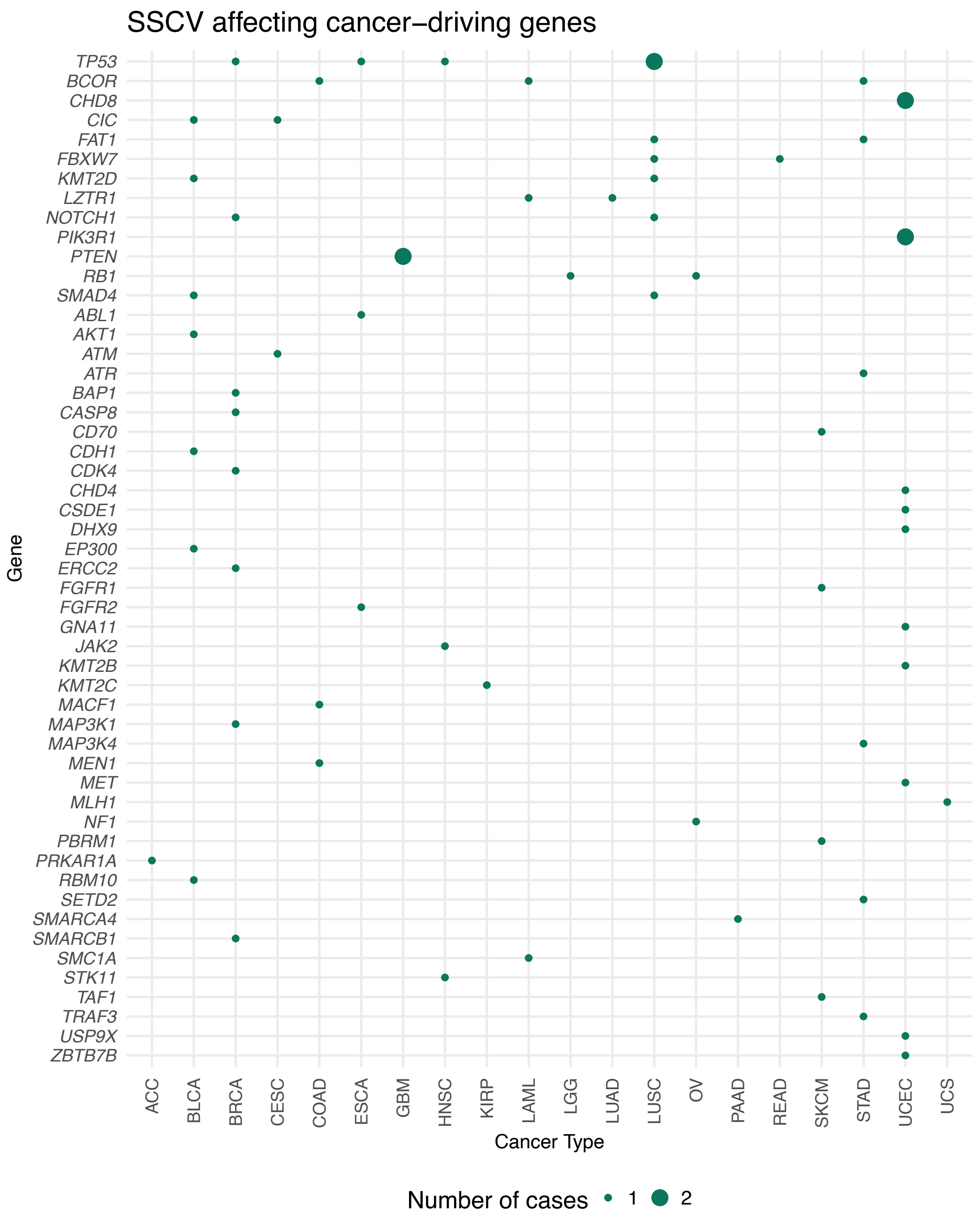


**Supplementary Figure 6: Landscape of SSCVs in cancer-related genes across cancer types.** The list of cancer driver genes was adopted from Bailey et al., 2018. Variants with allele frequencies in gnomAD greater than or equal to 0.0001 were excluded. The size of each point indicates the number of affected participants.

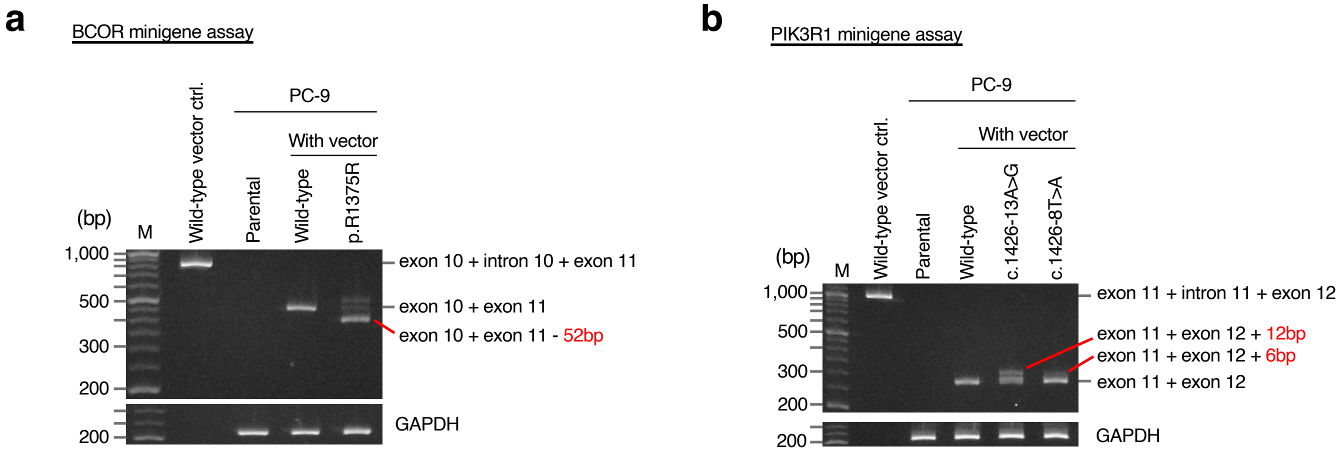


**Supplementary Figure 7: Validation of SSCVs affecting cancer-related genes via minigene-assay.** (a) Images of PCR amplicons targeting minigene-derived sequences, but not the endogenous sequences of *BCOR*. The control minigene vector itself shows the band spanning from exon 10, intron 10, to exon 11. PC-9 cells expressing *BCOR* p.R1375W minigene show alternative splicing and the band of loading control GAPDH, which is ubiquitously expressed in human cells. (b) Images of the *PIK3R1* minigene assay, conducted in the same manner. The *PIK3R1* mutants (c.1426-13A>G and c.1426-8T>A) also show alternative splicing.

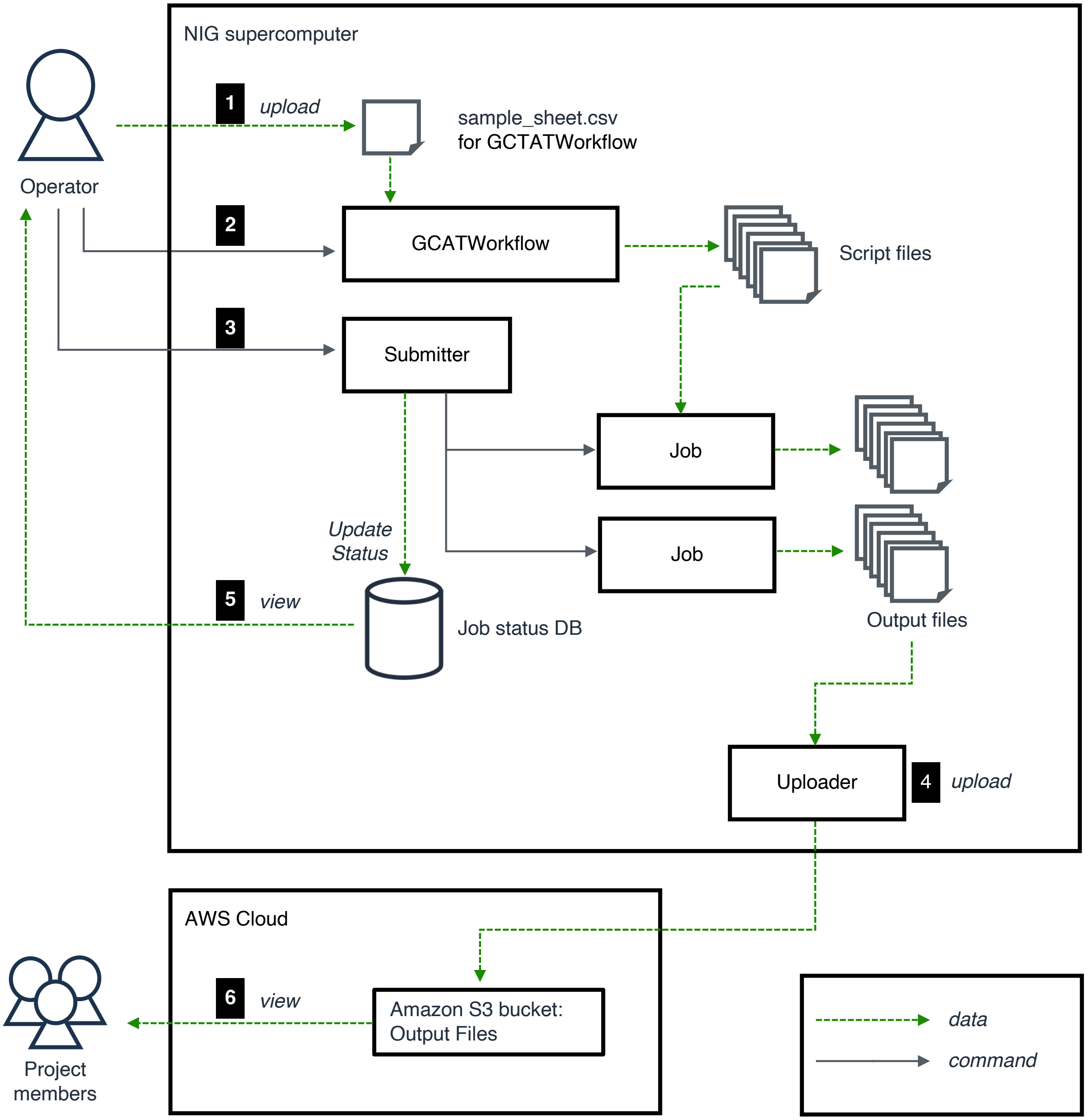


**Supplementary Figure 8: A workflow for detecting SSCVs using the NIG supercomputer.** We developed a snakemake based pipeline, GCATWorkflow (https://github.com/ncc-gap/GCATWorkflow), to execute downloading FASTQ files, alignment to the reference genome, and SSCV detection in order. Also, two background jobs (uploader and notifier) upload the output to Amazon S3 and notify the success or failure to Slack, respectively.

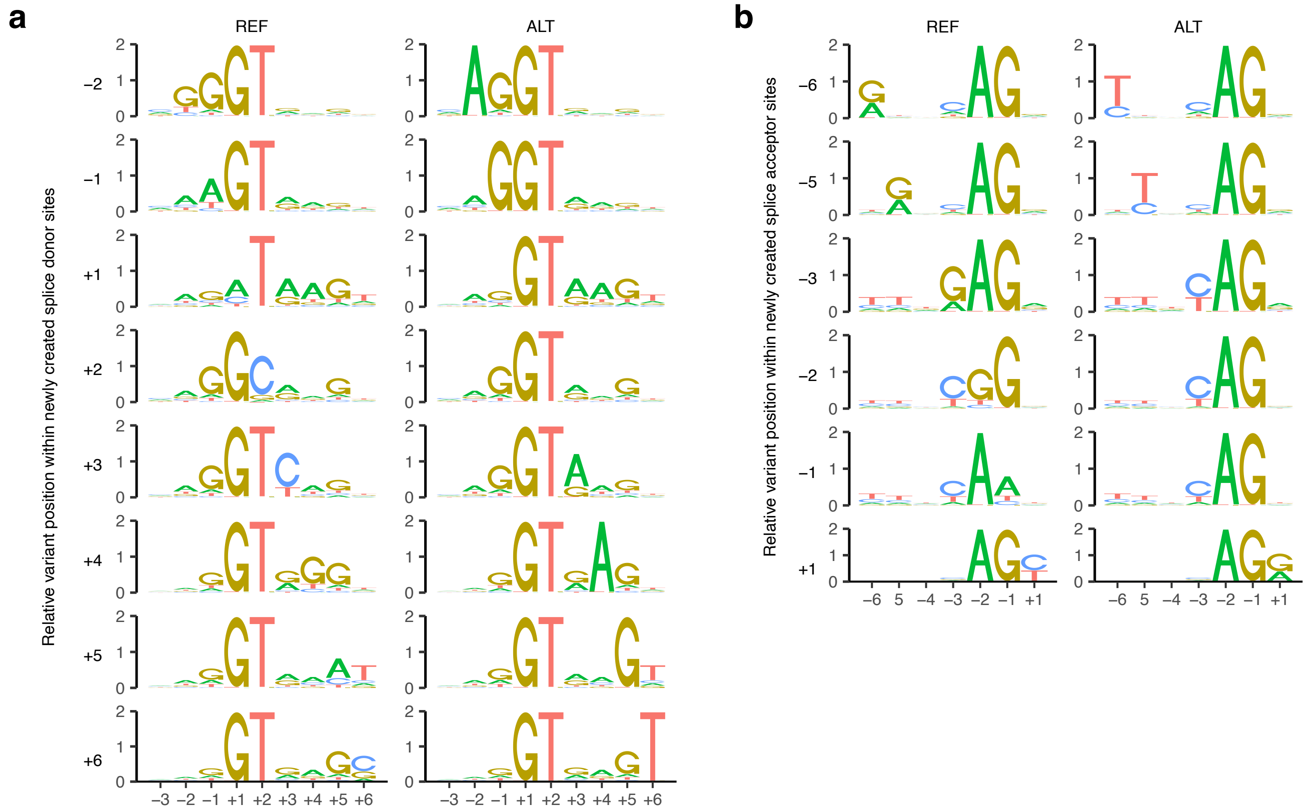


**Supplementary Figure 9: Sequence motifs of novel splice-sites by SSCVs.** (a, b) Novel donor (a) and acceptor (b) sites with (ALT) and without (REF) the SSCVs are categorized based on their relative variant position. Based on the relative position of the SSCVs to the newly generated splice-sites, newly created splice donor sequences were divided into left- and right-handed categories, with those exhibiting greater entropy positioned within the exon and the intron, respectively.

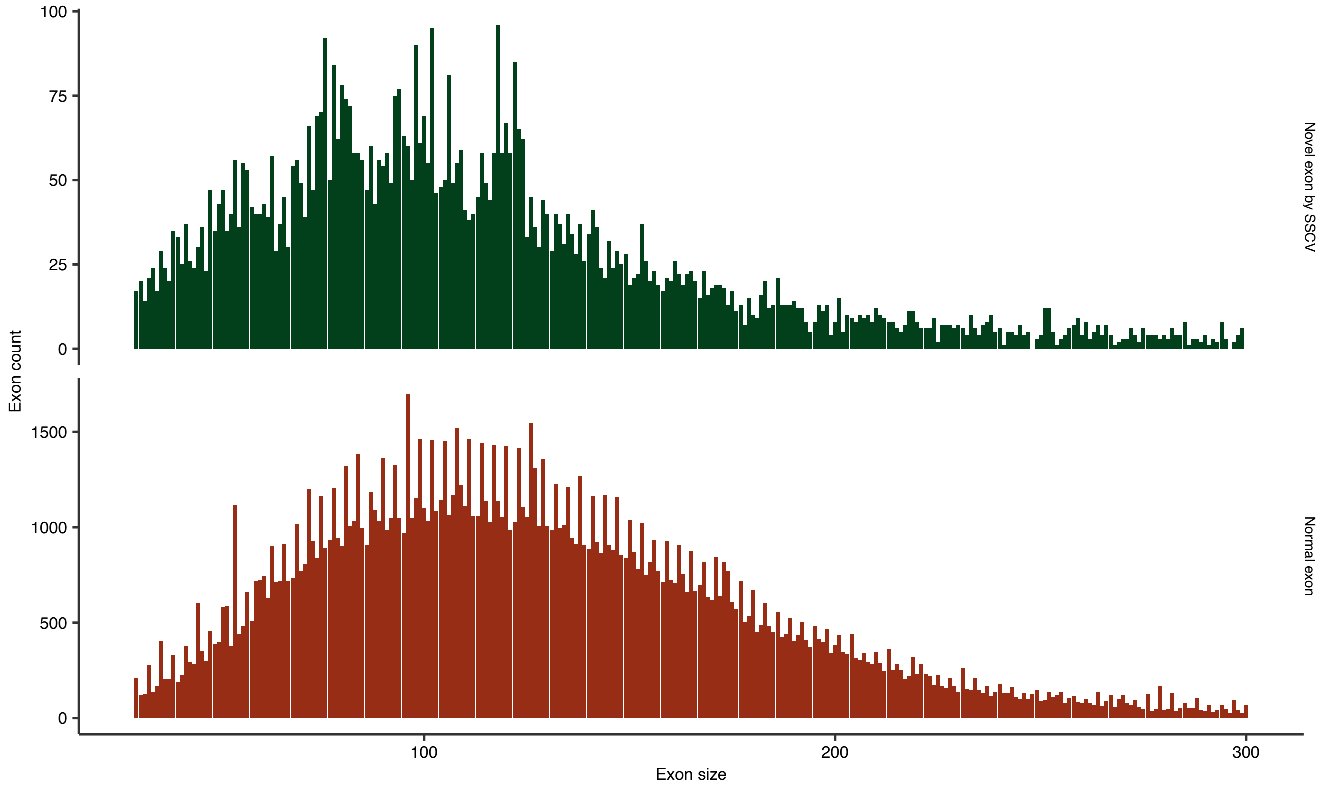


**Supplementary Figure 10: Histogram presenting the sizes of novel exons generated by SSCVs, which are predicted to lead to cryptic exon inclusion.** For comparative reference, the sizes of normal exons (excluding the first and last) in MANE transcripts are also depicted. A higher prevalence of changes in multiples of three is observed in these normal exons. This trend may be attributable to an evolutionary preference for multiples of three in exons, enhancing their resilience to exon-skipping events. Conversely, the frequent occurrence of new exon sizes being multiples of three caused by SSCVs might result from multiples of three being less susceptible to NMD, which could lead to a potential bias in their detection.

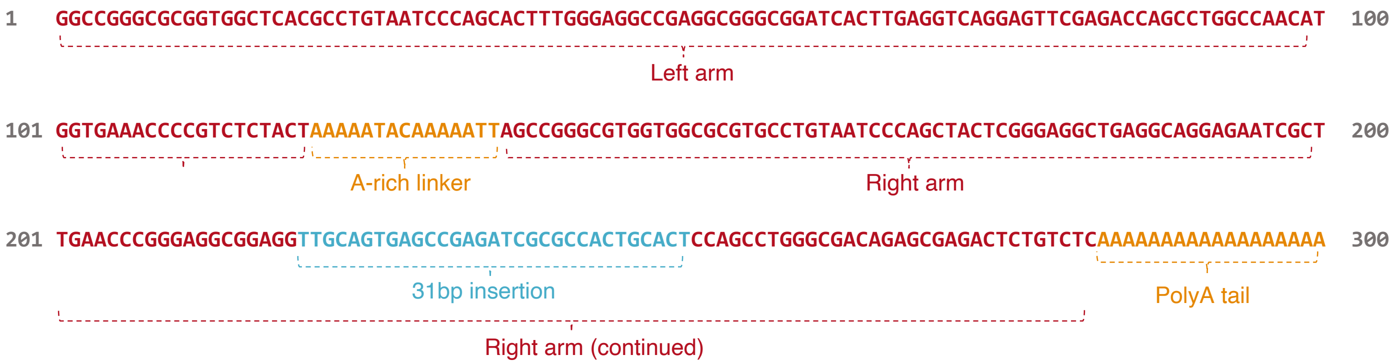


**Supplementary Figure 11: Reference Alu sequence used in this study.** This sequence was extracted from the study conducted by Funakoshi et al. The regions of the left arm, right arm, the 31 bp insertion sequence, A-rich linker, and the poly-A tail are distinctively colored.

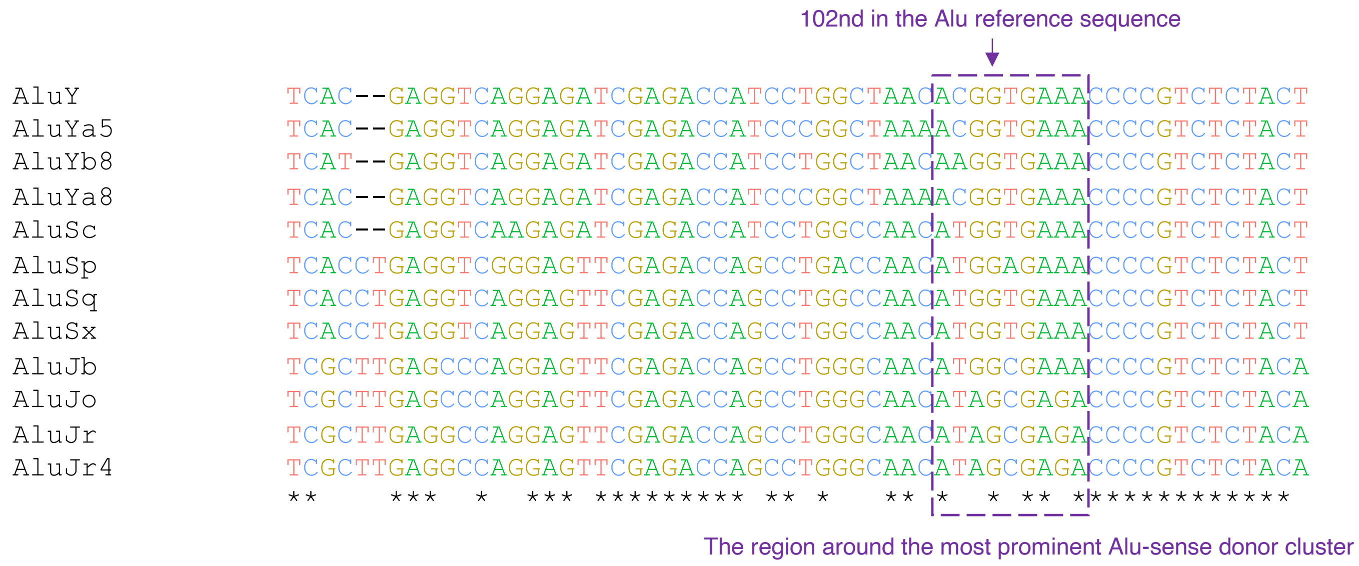


**Supplementary Figure 12: The results of a multiple alignment of sequences from several Alu subfamilies.** Here, particular focus is given to the most prominent Alu-sense donor hotspot (102nd). In AluJ, the first two letters of the intron sequence corresponding to the primary novel SS are GC, which could be converted to GT by SSCVs. In other Alu subfamilies, these letters are already GT; however, there may be mechanisms in place that inhibit splicing (e.g., the 5th intronic sites are mostly A).

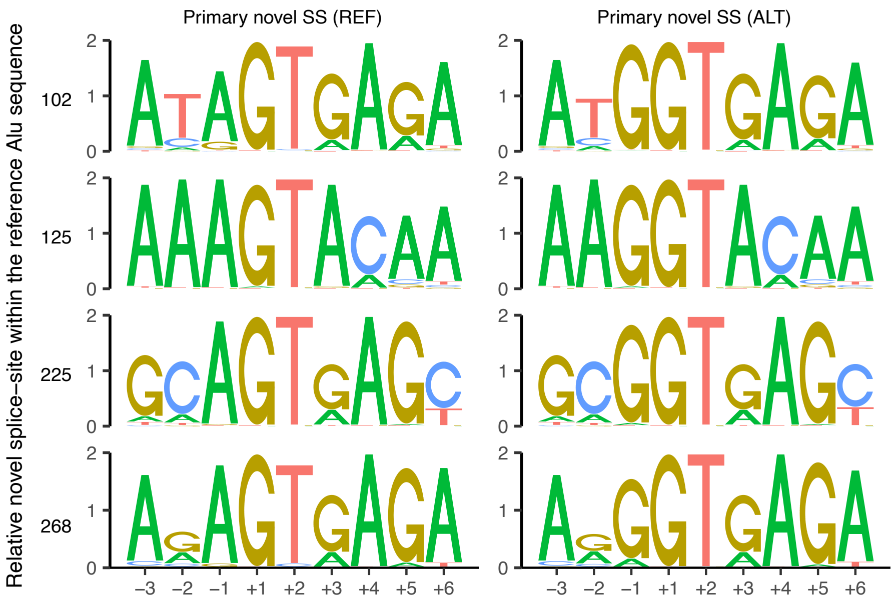


**Supplementary Figure 13: Sequence motifs of novel donor sites created by SSCVs within Alu sequences in sense direction.** For each relative position of primary novel SS within the reference Alu sequence (restricted to Alu-sense donor clusters along with the 125th position), primary novel SSs directly formed by SSCVs for both sequences before (REF) and after the mutation (ALT), are collected and their sequence motifs are evaluated.

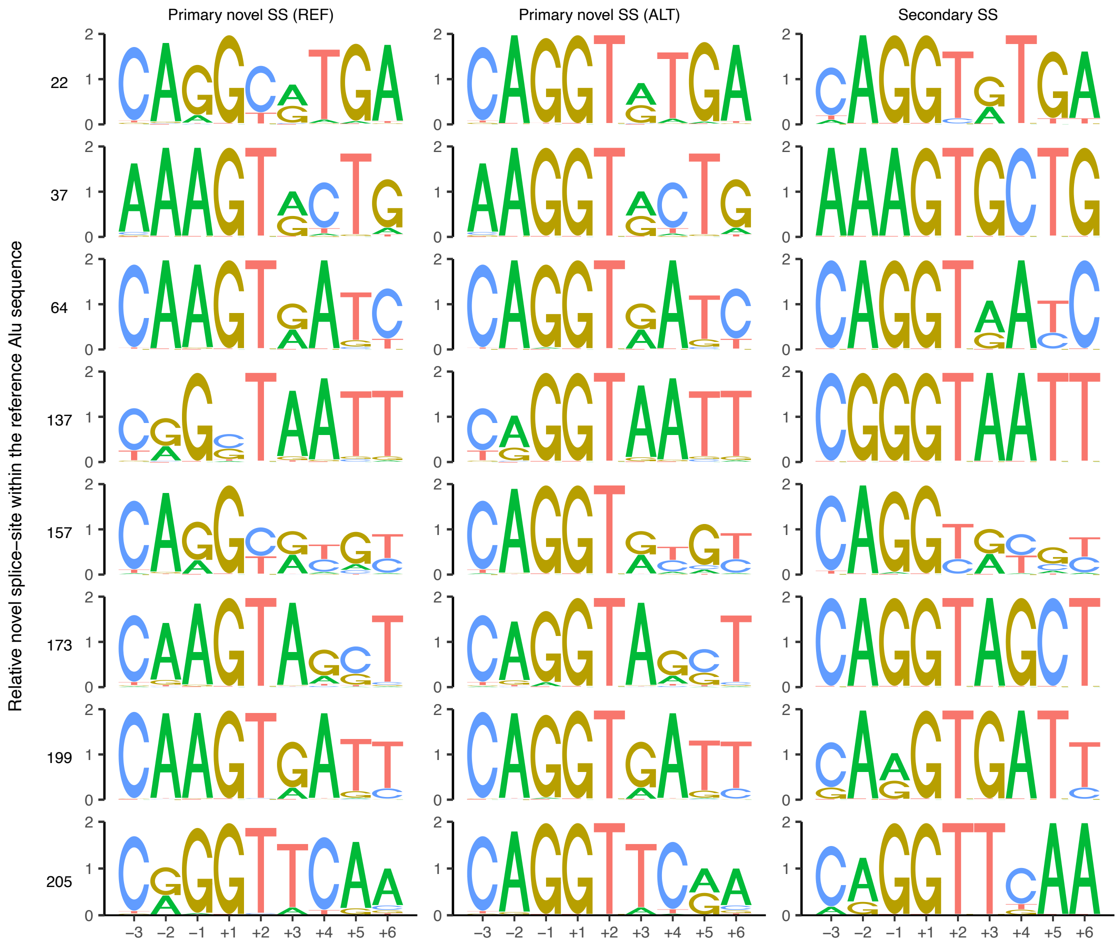


**Supplementary Figure 14: Sequence motifs of novel donor sites directly or indirectly created by SSCVs within Alu sequences in antisense direction.** For each relative position of primary novel SS within the reference Alu sequence (restricted to Alu-antisense donor clusters), primary novel SSs directly formed by SSCVs for both sequences before (REF) and after the mutation (ALT), as well as those secondarily formed through acceptor creation (Secondary SS), are collected and their sequence motifs are evaluated.

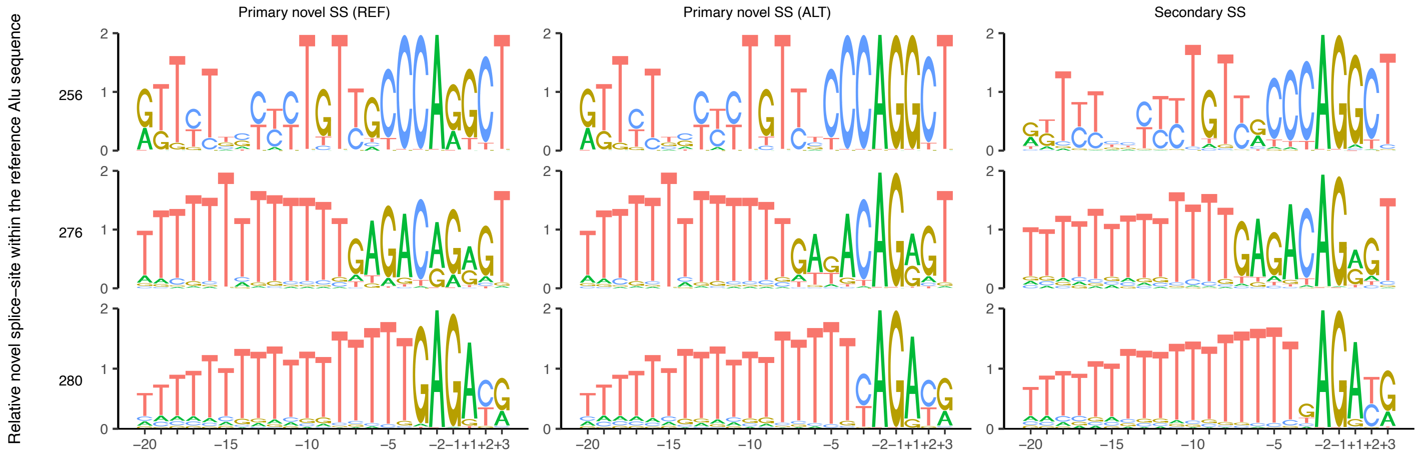


**Supplementary Figure 15: Sequence motifs of novel acceptor sites directly or indirectly created by SSCVs within Alu sequences in antisense direction.** For each relative position of primary novel SS within the reference Alu sequence (restricted to Alu-antisense acceptor clusters as well as the 256th position), primary novel SSs directly formed by SSCVs for both sequences before (REF) and after the mutation (ALT), as well as those secondarily formed through donor creation (Secondary SS), are collected and their sequence motifs are evaluated.

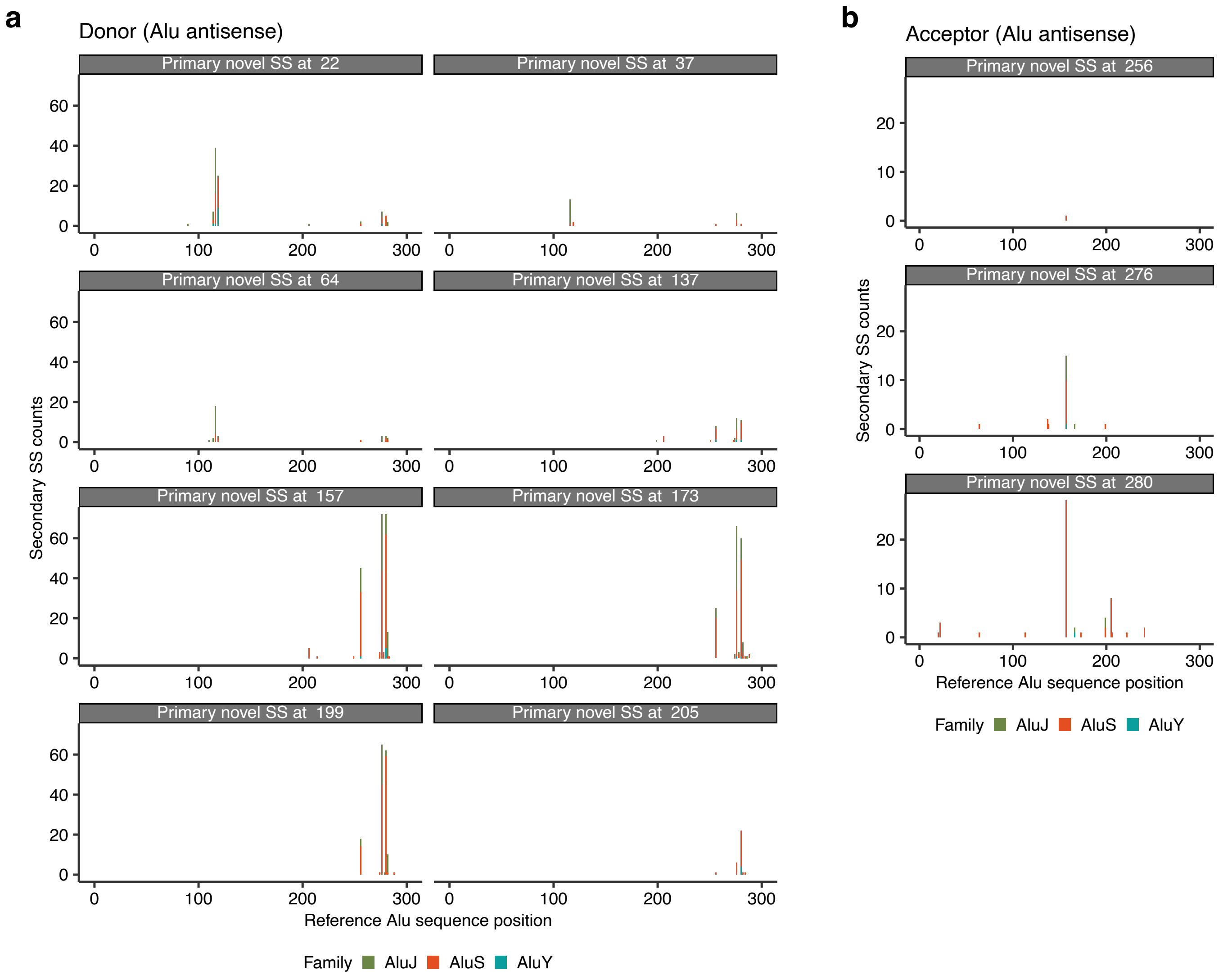


**Supplementary Figure 16: Distribution of secondary SS in Alu exonization by the SSCVs.** (a, b) The number of corresponding secondary SS at each position, mapped to reference Alu sequence coordinates, for SSCVs classified based on their primary novel SS (restricted to Alu-antisense donor (a) and acceptor (b) clusters) are shown. The counts are stratified by Alu family (AluJ, AluS, and AluY).

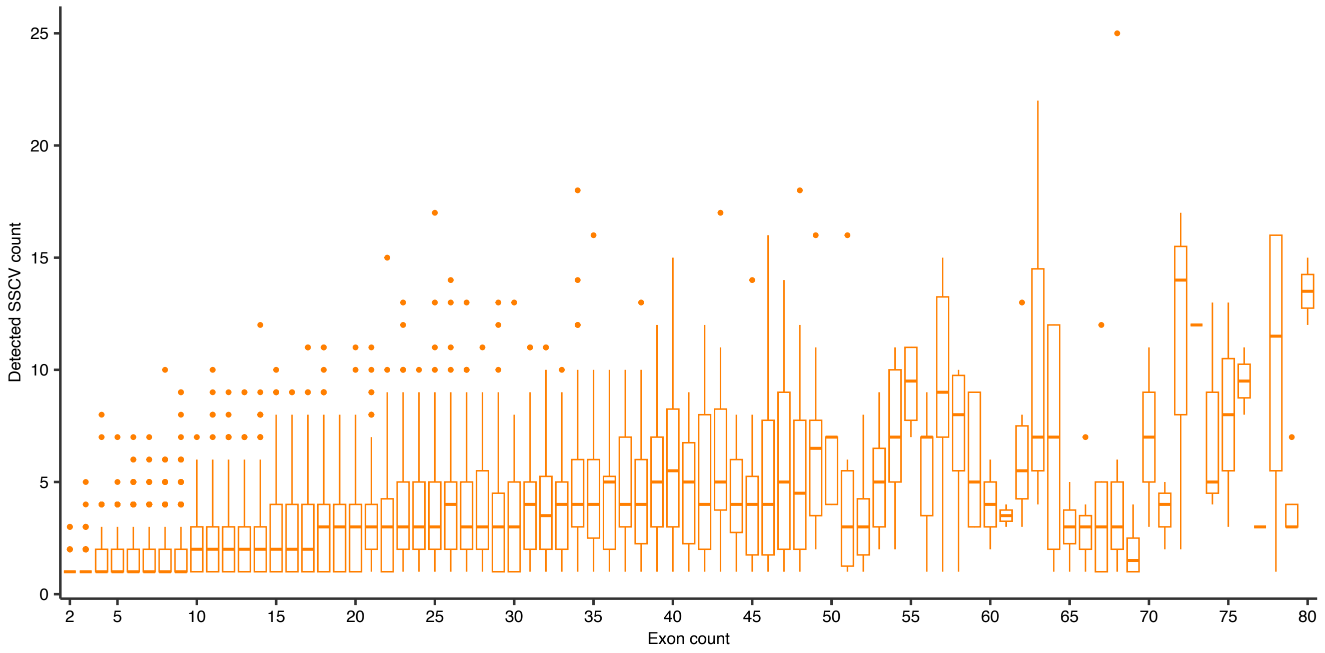


**Supplementary Figure 17: Boxplot illustrating the relationship between the number of exons and detected SSCV counts in genes.** Each box represents the range of SSCV counts observed in genes with a specific number of exons.

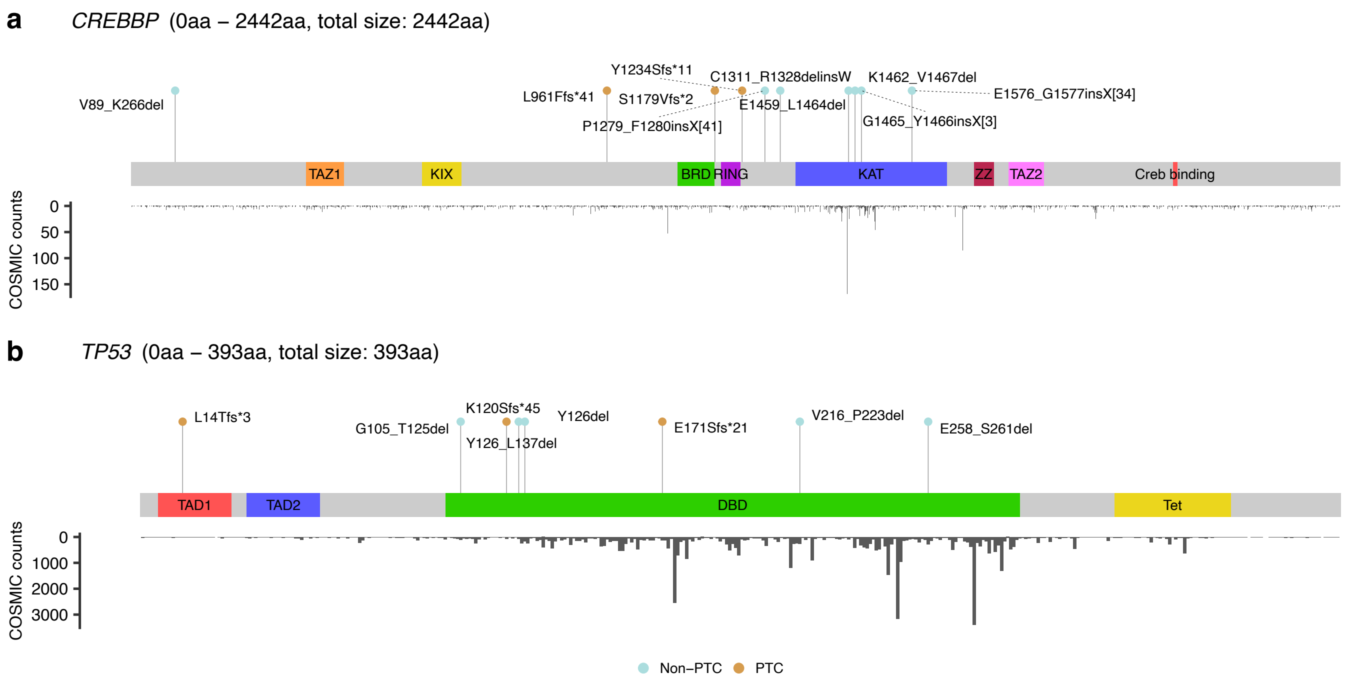


**Supplementary Figure 18: Lollipop plots of the detected SSCVs affecting *CREBBP* (a), *TP53* (b) across their entire amino acid regions.** These plots display labels denoting protein changes based on transcriptional consequences (using HGVS notation), and points are colored according to PTC generation (light blue for non-PTC, brown for PTC). Each plot is accompanied by the counts of somatic mutations at each amino acid position, as recorded in COSMIC.Unlike Figure 5c and 5d, this presentation displays all amino acid regions. See also Figure 5b-d and Supplementary Figure 19.

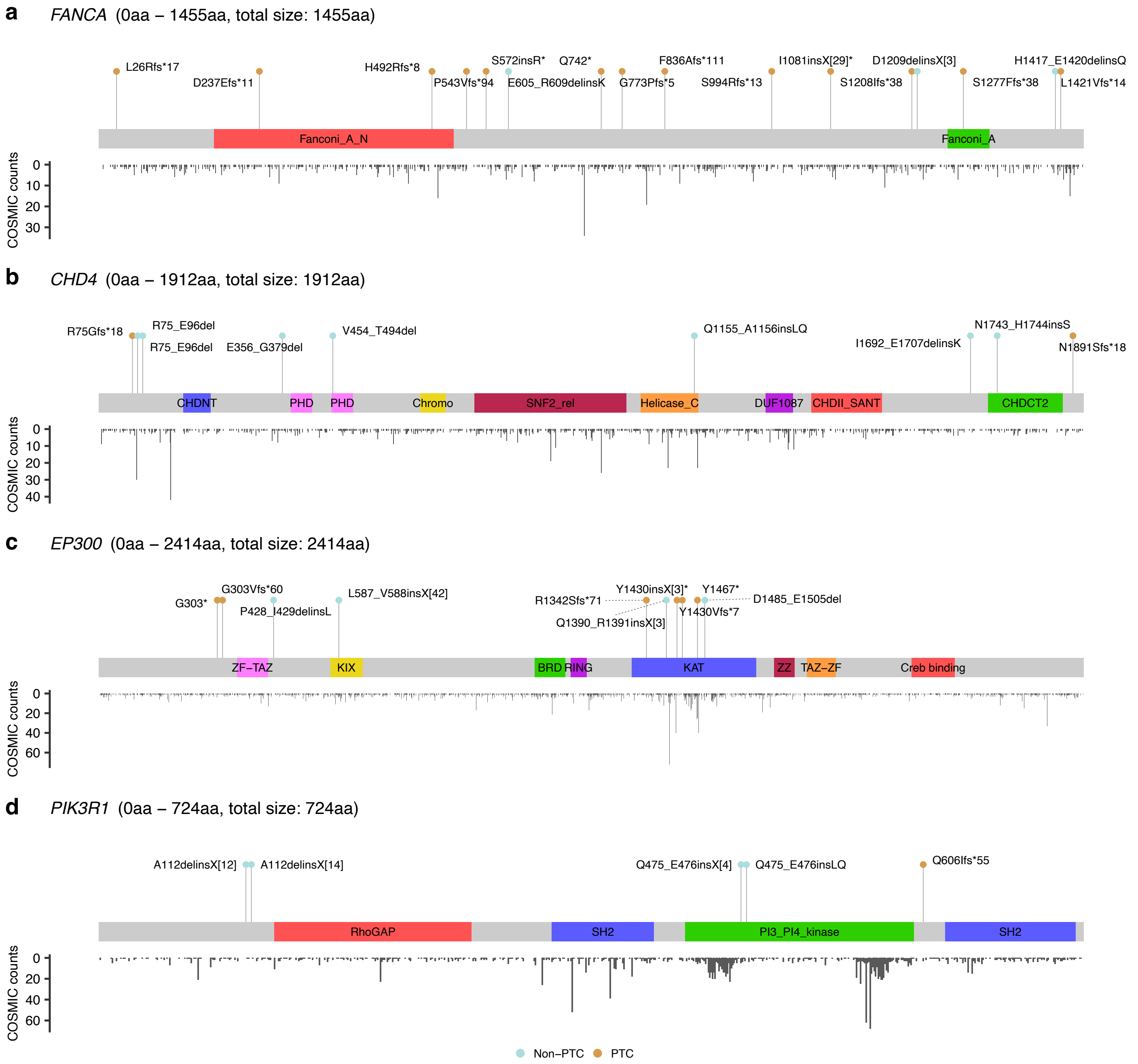


**Supplementary Figure 19: Lollipop plots of the detected SSCVs affecting *FANCA* (a), *CHD4* (b), *EP300* (c), *PIK3R1* (d).** These plots display labels denoting protein changes based on transcriptional consequences (using HGVS notation), and points are colored according to PTC generation (light blue for non-PTC, brown for PTC). Each plot is accompanied by the counts of somatic mutations at each amino acid position, as recorded in COSMIC. See also Figure 5b-d and Supplementary Figure 18.

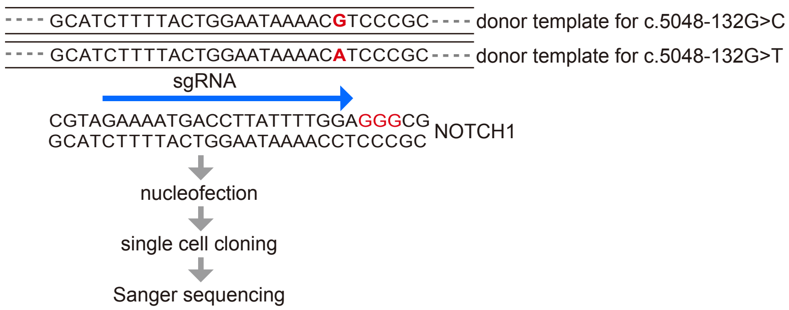


**Supplementary Figure 20: Workflow of *NOTCH1* mutant cell line generation using CRISPR-Cas9.** PC-9 cells were infected via nucleofection with sgRNA and donor templates designed to incorporate the c.5048-132G>C and c.5048-132G>T mutations. These *NOTCH1* mutations in single clones were confirmed by Sanger sequencing.

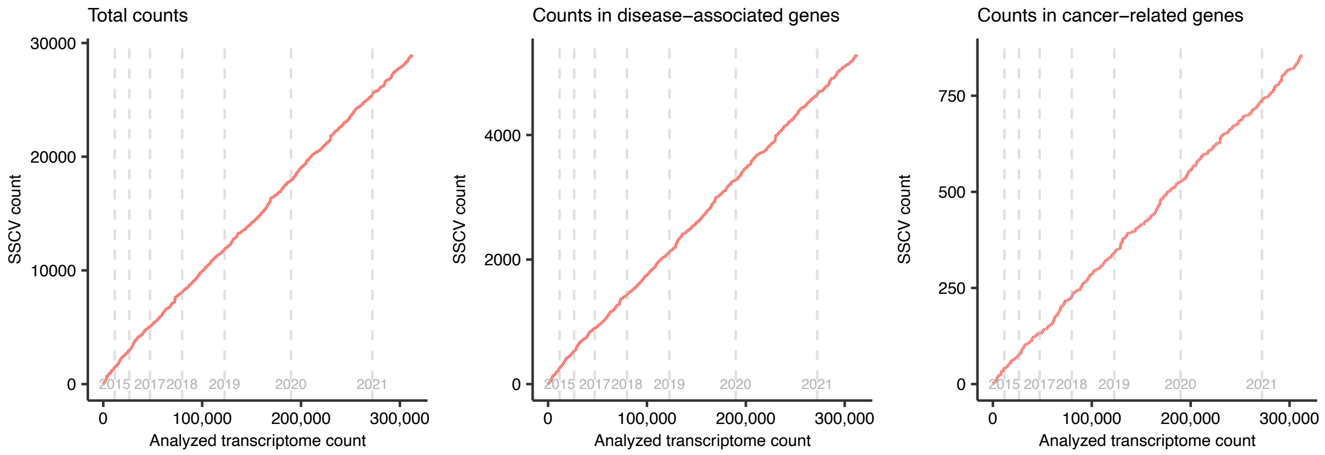


**Supplementary Figure 21: The saturation analysis of SSCVs via using Sequencing Read Archive.** Transcriptome data were sorted by registration date, and the cumulative number of SSCVs is shown. This includes all SSCVs (a), those affecting disease-associated genes (b), and cancer-related genes (c), plotted against the number of analyzed transcriptomes. Gray vertical lines denote the changes in registration years.
